# The study on the impact of indoleacetic acid on enhancing the ability of the rumen’s original microecology to degrade aflatoxin B1

**DOI:** 10.1101/2024.08.02.606440

**Authors:** Jiajin Sun, Zhonghao Wang, Xinyu Yan, Yuqi Zhao, Li Tan, Xuning Miao, Rong Zhao, Wenjie Huo, Lei Chen, Qinghong Li, Qiang Liu, Cong Wang, Gang Guo

**Author notes:** Correspondence: Gang Guo. First authorship: These authors share firstauthorship (Jiajin Sun, Zhonghao Wang).

## Abstract

Aflatoxin B1 (AFB1) poses a serious threat to the health of cattle and is strictly monitored in their diets. Rumen microorganisms in cattle have the ability to degrade AFB1, but this capability is limited. Enhancing the degradation capacity of the original rumen microecology for AFB1 is a novel approach. From a molecular perspective,indoleacetic-3-acid (IAA) promotes the expression of cytochromes, which can improve the degradation of AFB1. Therefore, this study aims to investigate the impact of different concentrations of IAA on the degradation of AFB1 by the rumen microecology. Experiments used rumen fluid from three adult cows as donors, and the cows were all fed the same total mixed ration. Rumen fluid was collected from these three cannulated cows before morning feeding to prepare in vitro fermentation fluid. The experiments used a completely randomized design, with each treatment repeated four times. The results showed that as the fermentation time increased, the content of AFB1 gradually decreased, with a degradation rate of up to 75.73% after 24 h. AFB1 altered the rumen fermentation pattern, with a significant reduction in the content of acetic acid (*P*<0.05) and a significant decrease in the acetic acid to propionic acid ratio (*P*<0.05). It also affected the rumen microecology, causing a significant reduction in the abundance of *Ruminococcus amylophilus, Prevotella ruminicola*, and *Fusobacterium succinogenes* (*P*<0.05). In addition, this study found that with the increase in the amount of IAA added, the content of AFB1 in the rumen gradually decreased. IAA enhances the degradation capacity of the original rumen microecology for AFB1, and the addition of IAA alleviates the impact of AFB1 on *Ruminococcus amylophilus, Prevotella ruminicola*, and *Fusobacterium succinogenes* in the rumen (*P*<0.05). Moreover, the addition of IAA can promote the stability of the rumen microecology, with significantly higher acetic acid and acetic acid to propionic acid ratios in the fermentation fluid compared to the non-added group (*P*<0.05). In summary, the addition of IAA can improve the degradation capacity of the rumen microecology for AFB1, providing a new solution for alleviating the impact of AFB1 on animal health.

## 1 Introduction

Mycotoxins are toxic secondary metabolites produced by molds during their growth process and are widely present in grains and animal feed [1]. Among them, aflatoxins are considered to be important class I carcinogens, posing a serious threat to the health of humans and animals [2,3]. Therefore, controlling the content of aflatoxin B1 (AFB1) in animal diets is particularly important. In recent years, silage, corn, and its by-products have been widely used as the main feed materials for cattle, but frequent occurrences of AFB1 exceeding standards have led to serious waste of feed resources and economic losses [4,5]. Studies have shown that rumen microorganisms have the ability to degrade AFB1, but this ability is limited. This is mainly achieved through the following mechanisms: Biotransformation: Certain rumen microorganisms can convert AFB1 into metabolites with lower toxicity or non-toxic. Adsorption: Metabolites produced by some microorganisms or cell surface structures may have the ability to adsorb AFB1, thereby reducing its absorption in the body. Competitive inhibition: Rumen microorganisms may compete with AFB1 for absorption sites, reducing its absorption in the intestine. Enzymatic reactions: Specific microbial enzymes may catalyze the breakdown of AFB1, producing harmless or low-toxic metabolites [6,7]. However, the ability of rumen microorganisms to degrade AFB1 is limited, which may be affected by the following factors: Types and quantities of microorganisms: Different types of rumen microorganisms have different degradation capabilities for AFB1, and the composition and quantity of microbial communities can affect the efficiency of degradation. Feed components: Some components in the feed may affect the activity of microorganisms or their interaction with AFB1. Therefore, finding a plant extract additive to enhance the degradation capacity of the original rumen microecology for AFB1 is a new approach for degradation.

Fresh green forage contains some extracts and other features that are non-toxic and environmentally friendly, which will become a potential resource library for screening degradative toxins. Indoleacetic-3-acid (IAA), as a regulatory factor in plants, has various special functions in microbial metabolism [8,9], and has a certain regulatory effect on the synthesis of tryptophan and cytochromes by rumen microorganisms [10]. At the same time, IAA, as an activator of the cellular aryl hydrocarbon receptor (AhR), activates the expression of cytochrome genes such as CYP1A1, CYP1A2, and CYP1B1 in cells, playing a key role in the physiological functions and immune responses of cells. In addition, cytochrome P450 enzymes have been found in different organisms and are involved in the biotransformation of AFB1 [11]. CYP450 enzymes are a group of mixed-function oxidases involved in the biotransformation of endogenous compounds (such as bile acids, prostaglandins, steroids, fatty acids, etc.) and exogenous compounds (drugs, carcinogens, pro-mutagens), and are present in most organisms. In addition, IAA is involved in the metabolism of various beneficial bacteria [12,13] and promotes the growth of probiotics [14]. Probiotics have shown significant potential in degrading AFB1, with diverse mechanisms of action, including biodegradation, adsorption, competitive exclusion, regulation of the intestinal environment, enhancement of host immune response, gene regulation, and synergistic action with other detoxification enzymes [15,16]. These mechanisms work together to reduce the toxicity of AFB1, decrease its absorption in the body, and improve the host’s clearance ability [17,18]. However, there are few reports on the alleviation of the negative impact of AFB1 on rumen fermentation by adding IAA. Therefore, the coordination of rumen microorganisms by IAA to achieve the degradation of AFB1 in the rumen requires further study.

In summary, as a potential rumen regulatory substance, whether IAA can promote the degradation of AFB1 in the rumen is not yet clear. Therefore, this study investigates the impact of IAA on the degradation of AFB1 in the rumen by adding IAA, elucidates the impact of IAA on the degradation of AFB1 in the rumen, and promotes the research on the synergistic degradation of AFB1 by harmless chemicals and intestinal microorganisms. This work is of great significance for the degradation of mycotoxins in feed and raw materials. It can not only improve the safety of animal feed but also reduce the waste of grain, which has an important impact on the long-term and stable development of China’s animal husbandry and agriculture.

## 2 Materials and Methods

### 2.1. Experimental design

1. Alfalfa silage and starch, each 0.2 g, were weighed and mixed in nylon bags and placed in fermentation bottles in advance. 60 mL of rumen fluid and 40 μL of AFB1 standard solution (10 μg/mL) were added to the bottles, setting the AFB1 concentration to 1 mg/kg. The rumen fluid fermentation bottles containing 1 mg/kg AFB1 were divided into 8 groups, with 4 replicates each, and placed in a 39°C constant temperature shaker for in vitro fermentation at 0, 1, 2, 3, 4, 5, 24, and 48 h. After the fermentation, pre-column derivatization [19] was carried out to detect AFB1 content and rumen fermentation indicators.
2. Alfalfa silage and starch, each 0.2 g, were weighed and mixed in nylon bags and placed in fermentation bottles in advance. 60 mL of rumen fluid and 40 μL of AFB1 standard solution (10 μg/mL) were added to the bottles, setting the AFB1 concentration to 1 mg/kg. The rumen fluid fermentation bottles containing 1 mg/kg AFB1 were supplemented with IAA standard solution (600 μg/mL) in amounts of 0, 10 μL, 100 μL, 1 mL, and 5 mL, setting the IAA concentrations in the 5 fermentation bottles to 0, 15 mg/kg, 150 mg/kg, 1500 mg/kg, and 7500 mg/kg, respectively. Each group had 4 replicates and was placed in a 39°C constant temperature shaker for in vitro fermentation for 24 h. After pre-column derivatization [19], the AFB1 content and rumen fermentation indicators were detected.

### 2.2 Experimental materials and animals

In this experiment, three healthy adult cows with similar body conditions and ages were selected as rumen fluid donors. IAA (Shanghai Macklin Biochemical CO, LTD), Aflatoxin B1 (Stanford Analytical Chemicals Inc).

### 2.3 Collection and in vitro cultivation method of rumen fluid

Before sampling, the fermentation bottles were preheated in a 39°C constant temperature incubator. Before the morning feeding (07:00), rumen fluid was collected from the three Holstein cows through the oral cavity. The mixed rumen fluid was filtered through four layers of gauze and temporarily stored in a thermos preheated to 39°C and supplied with carbon dioxide (CO_2_), maintaining an anaerobic environment. The rumen fluid was mixed with artificial saliva in a 1:1 ratio as described by Longland *et al.* [20] and continuously supplied with CO_2_. Alfalfa silage and starch were mixed in a 1:1 ratio and placed in nylon bags. The reaction mixtures and collected rumen fluid were evenly mixed, and each 60 mL was divided into individual fermentation bottles, sealed, and placed in a 39°C shaker for cultivation [21] (speed of 120 r/min). When the fermentation time reached the set value, it was immediately removed, stopped in an ice bath, and samples were taken for detection of various indicators.

### 2.4 Determination items and methods

After 24 h of in vitro fermentation, the fermentation fluid was cooled and the contents of AFB1, ammonia nitrogen (NH_3_-N), volatile fatty acids (VFA), and four types of cellulase were measured.

The content of AFB1 was determined using an Agilent 1260 Infinity II high-performance liquid chromatograph (HPLC) with a C18 column, 4.6×250 nm, 5 μm. The mobile phase was acetonitrile-water (20:80); column temperature: 40°C; mobile phase flow rate: 1.0 mL/min; injection volume: 20 μL; excitation wavelength 360 nm, emission wavelength 440 nm; detection time 20 min.

The degradation rate of AFB1 = (AFB1 content in the blank control - AFB1 content in the experimental group) / AFB1 content in the blank control × 100%.

The content of ammonia nitrogen was determined using the phenol-hypochlorous acid colorimetric method [22].

pH was measured using a pH meter (LE438, Mettler Toledo Instruments Co., Ltd.).

The content of volatile fatty acids (VFA) was determined using a high-performance gas chromatograph (GC-TRACE 1300, column model 30 m×0.25 mm×0.25 μm) [23].

The determination of the four types of cellulase was carried out according to the method described by Agarwal *et al.* [24].

Microbial DNA was extracted using the bead-CTAB method, and real-time fluorescent quantitative PCR (ABI StepOnePlus, USA) was performed for RT-PCR. [25] (Primers are in the attachment)

### 2.5 Data processing and statistical analysis

Experimental data were recorded using Excel, and data analysis was performed using SPSS software, with paired sample T-tests and repeated measures models. The level of statistical significance was set at *P*<0.05, and the level of highly significant difference was set at *P*<0.001. Figures were drawn using KingDraw, Prism 9, and Adobe Illustrator 2022 for graphic illustration.

## 3. Results

### 3.1. The impact of different fermentation times on the degradation rate of AFB1 in vitro fermentation fluid

Figure 1(a) reveals that during the in vitro fermentation process of alfalfa silage, the degradation efficiency of AFB1 shows a stable and significant increasing trend as the reaction time is extended. In the 0-24 hour reaction, the degradation rate rises rapidly; between 24-48 h, the rate of increase in degradation slows down. When the reaction reaches 48 h, the fermentation fluid’s degradation rate of AFB1 can reach 80.09%. Taking all factors into consideration, this experiment selects 24 h as the final fermentation time.

**Fig.1.**
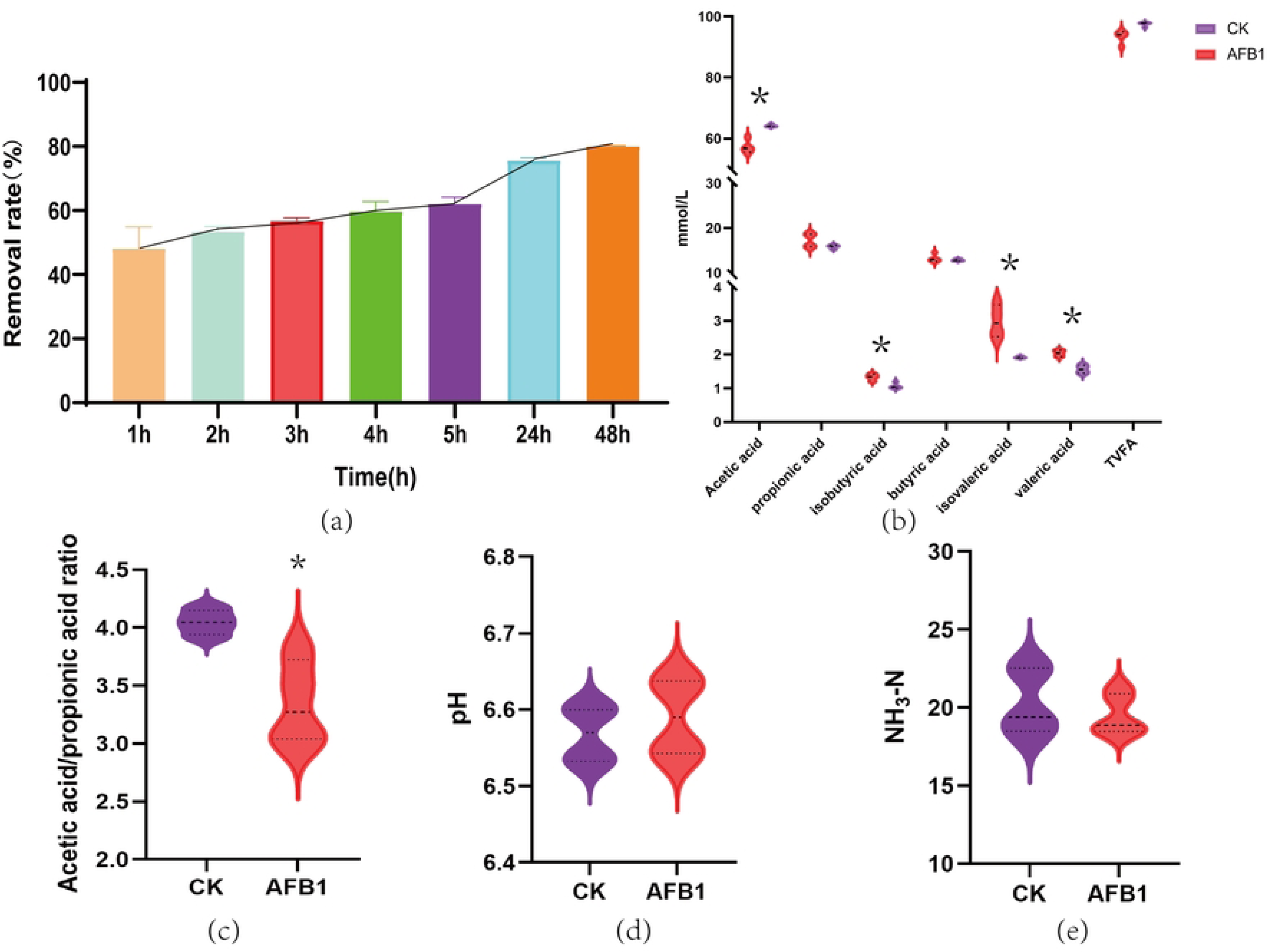
(a) The effect of different fermentation time on the degradation rate of AFB1 in in vitro fermentation broth. (b) volatile fatty acids, (c) Acetic acid/propionic acid ratio, (d) pH, (e) ammonia nitrogen production. Consistent with * means *P* < 0.05, * * means *P* < 0.001.

### 3.2 The impact of AFB1 on the production of volatile fatty acids, pH, and ammonia nitrogen in vitro fermentation fluid

Data from Figures 1(b, c, d, e) indicate that in the alfalfa silage in vitro fermentation fluid, compared to the control group, the addition of 1 mg/kg AFB1 significantly reduced the content of acetic acid (*P*<0.05); meanwhile, the content of propionic acid, isobutyric acid, valeric acid, and isovaleric acid significantly increased (*P*<0.05). Further analysis shows that the addition of 1 mg/kg AFB1 to the in vitro fermentation fluid led to a significant decrease in the acetic acid to propionic acid ratio (*P*<0.05). In addition, the addition of AFB1 did not significantly affect the pH value and ammonia nitrogen content of the in vitro fermentation fluid.

### 3.3 The impact of AFB1 on the activity of major fiber-degrading enzymes in vitro fermentation fluid

The impact of adding 1 mg/kg AFB1 on the activity of major cellulose-degrading enzymes in the rumen fermentation fluid is shown in Figure 2. It can be seen from Figure 2 that the addition of AFB1 has a highly significant effect on reducing the activity of xylanase after in vitro fermentation of alfalfa silage (*P*<0.05), while it does not significantly affect the activity of pectinase, carboxymethyl cellulase sodium, and β-glucosidase.

**Fig.2.**
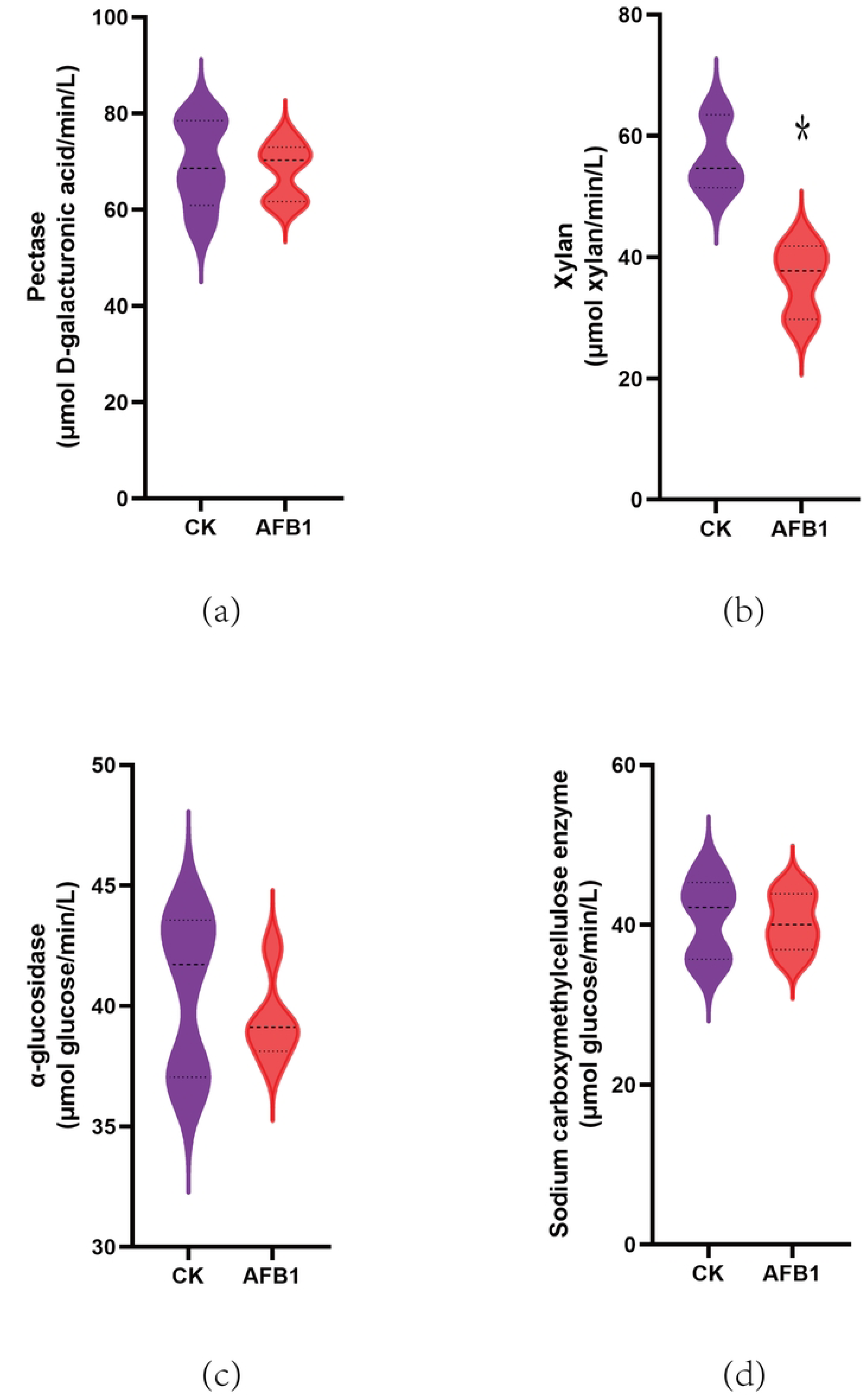
Effect of AFB1 on the activity of major fiber degrading enzymes in vitro fermentation broth, * indicates P < 0.05, * * indicates P < 0.001.

### 3.4 The impact of AFB1 on the microbial flora in vitro fermentation fluid

Data from Figure 3 reveal that after the addition of 1 mg/kg AFB1, the content of *Prevotella ruminicola* and *Fusobacterium succinogenes* significantly decreased (*P*<0.05), indicating that the presence of AFB1 inhibits the growth of these two types of bacteria. This result has potential implications for the balance of the microbial community within the rumen.

**Fig.3.**
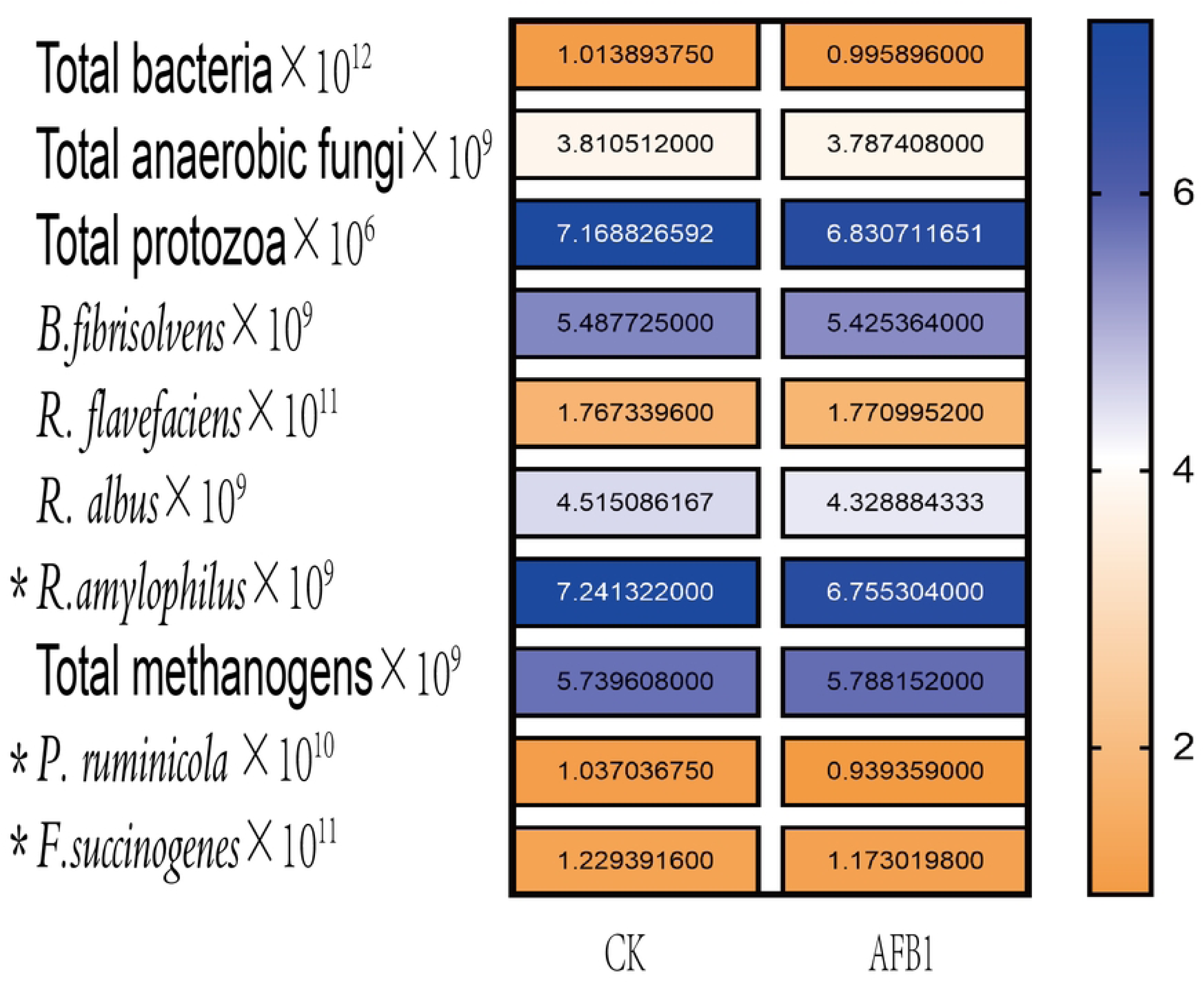
Effect of AFB1 on microbial flora in in vitro fermentation broth, * indicates P < 0.05, * * indicates P < 0.001.

### 3.5 The impact of different concentrations of IAA on the degradation rate of AFB1 in 24-hour in vitro fermentation fluid

Figure 4 illustrates that compared to the control group with only 1 mg/kg AFB1 added, there is a noticeable upward trend in the degradation rate of AFB1 in the alfalfa silage in vitro fermentation fluid as the amount of IAA increases. It can be observed that there is a certain positive correlation between the amount of IAA added and the degradation rate of AFB1 by the rumen fermentation fluid. As the addition of IAA gradually increases, the degradation effect on AFB1 by the rumen fermentation fluid is enhanced. When the IAA addition reaches 7500 mg/kg, the degradation rate of AFB1 by the fermentation fluid peaks at 75.1%.

**Fig.4.**
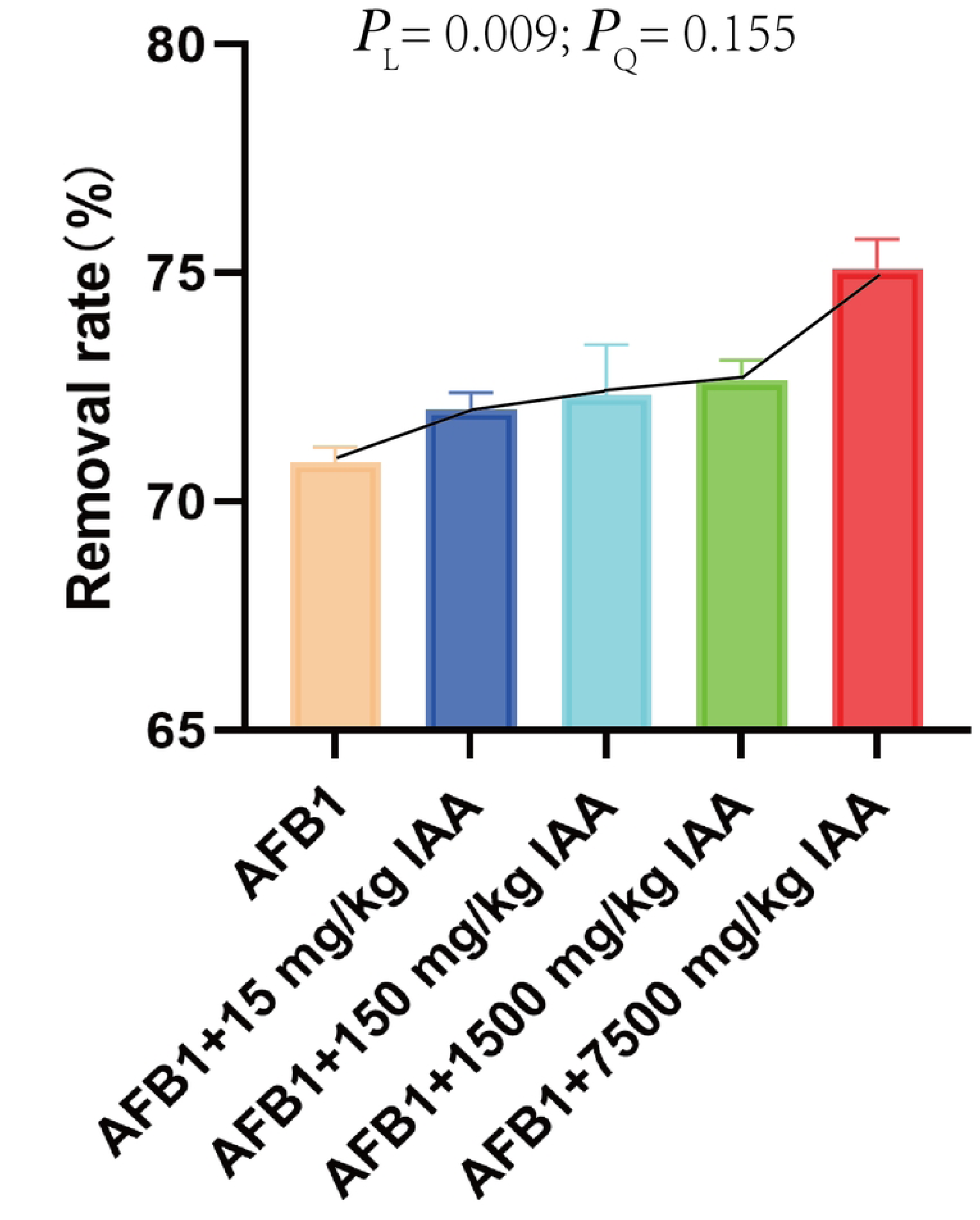
Effects of different concentrations of IAA on the degradation rate of AFB1 in 24 h fermentation broth in vitro. P_L_ represents a linear response, P_Q_ represents a quadratic linear response.

### 3.6 The impact of different concentrations of IAA on the production of volatile fatty acids, pH, and ammonia nitrogen in 24-hour in vitro fermentation fluid

Figure 5 indicates that compared to the control group with only 1 mg/kg AFB1 added, the addition of low and medium concentrations of IAA shows a positive correlation between the amount of IAA and the content of volatile acids in the fermentation fluid. Specifically, there is a significant increasing trend in the content of acetic acid, propionic acid, butyric acid, and total volatile fatty acids (TVFA) (linear effect, *P*=0.007, 0.028, 0.026, 0.006; quadratic effect, *P*=0.028, 0.021). Additionally, when the IAA addition is 150 mg/kg, there is also a significant increase in isovaleric acid in the fermentation fluid (*P*<0.05). Furthermore, when 1500 mg/kg IAA is added, the content of acetic acid, valeric acid, the acetic acid to propionic acid ratio, and TVFA in the fermentation fluid all significantly increase (*P*<0.05), while the content of propionic acid, isobutyric acid, butyric acid, and isovaleric acid significantly decrease (*P*<0.05). The addition of IAA did not significantly affect the pH and ammonia nitrogen content in the fermentation fluid.

**Fig.5.**
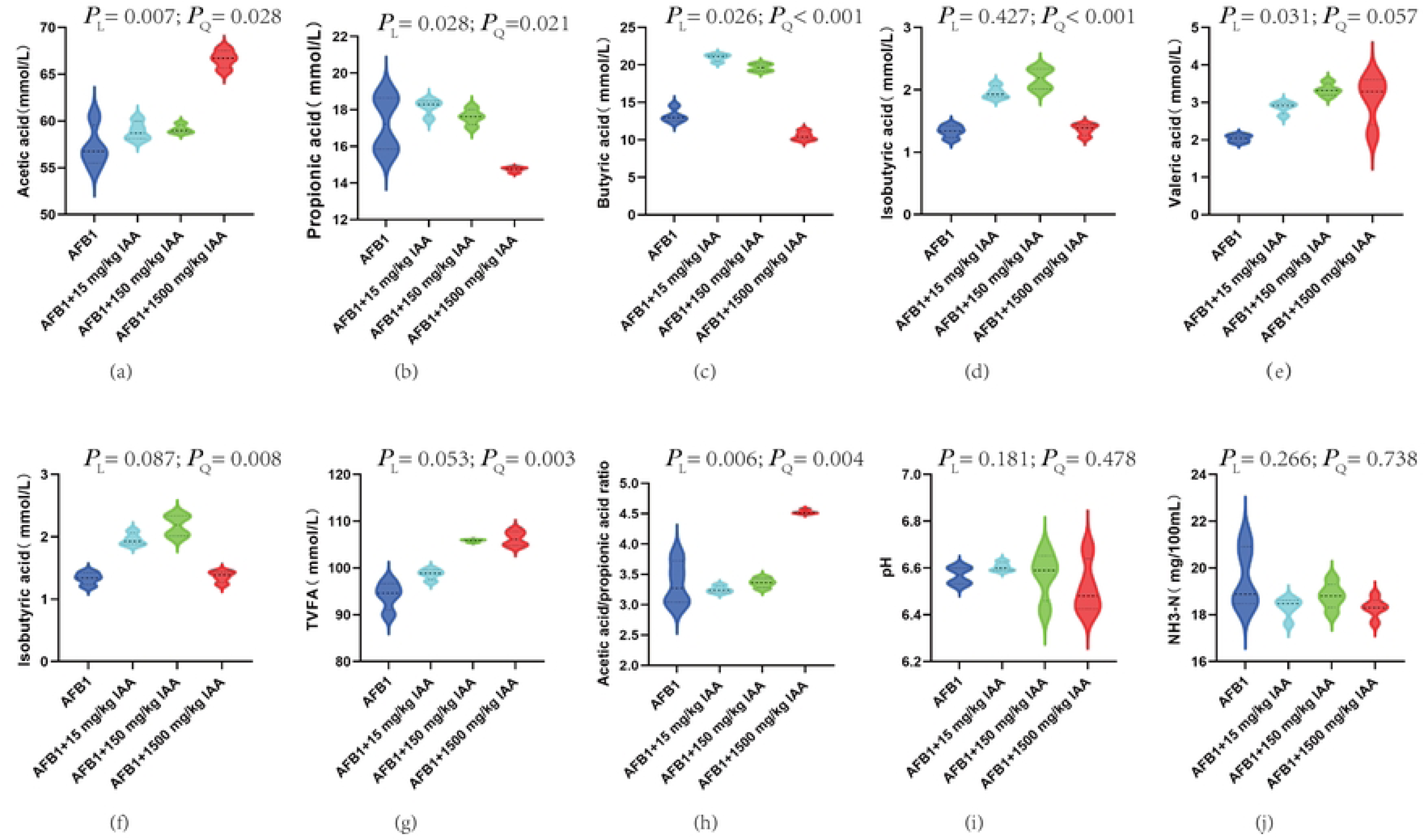
Effects of different concentrations of indole acetic acid on the production of volatile fatty acids, pH and ammonia nitrogen in 24 h in vitro fermentation broth. P_L_ represents a linear response, P_Q_ represents a quadratic linear response.

### 3.7 The impact of different concentrations of IAA on the activity of major fiber-degrading enzymes in 24-hour in vitro fermentation fluid

The impact of adding different concentrations of IAA on the activity of major fiber-degrading enzymes in the 24-hour in vitro fermentation fluid is shown in Figure 6. It can be seen that, compared to the control group with only 1 mg/kg AFB1 added, the addition of 15 and 150 mg/kg IAA shows an enhancing trend in the activity of pectinase, xylanase, and carboxymethyl cellulase sodium after the in vitro fermentation of alfalfa silage (where pectinase activity shows a linear effect, *P*=0.002; xylanase and carboxymethyl cellulase sodium activities show a quadratic effect, *P*=0.027, 0.005). In contrast, high concentrations of IAA exhibit a certain inhibitory effect on the activity of pectinase, xylanase, and carboxymethyl cellulase sodium.

**Fig.6.**
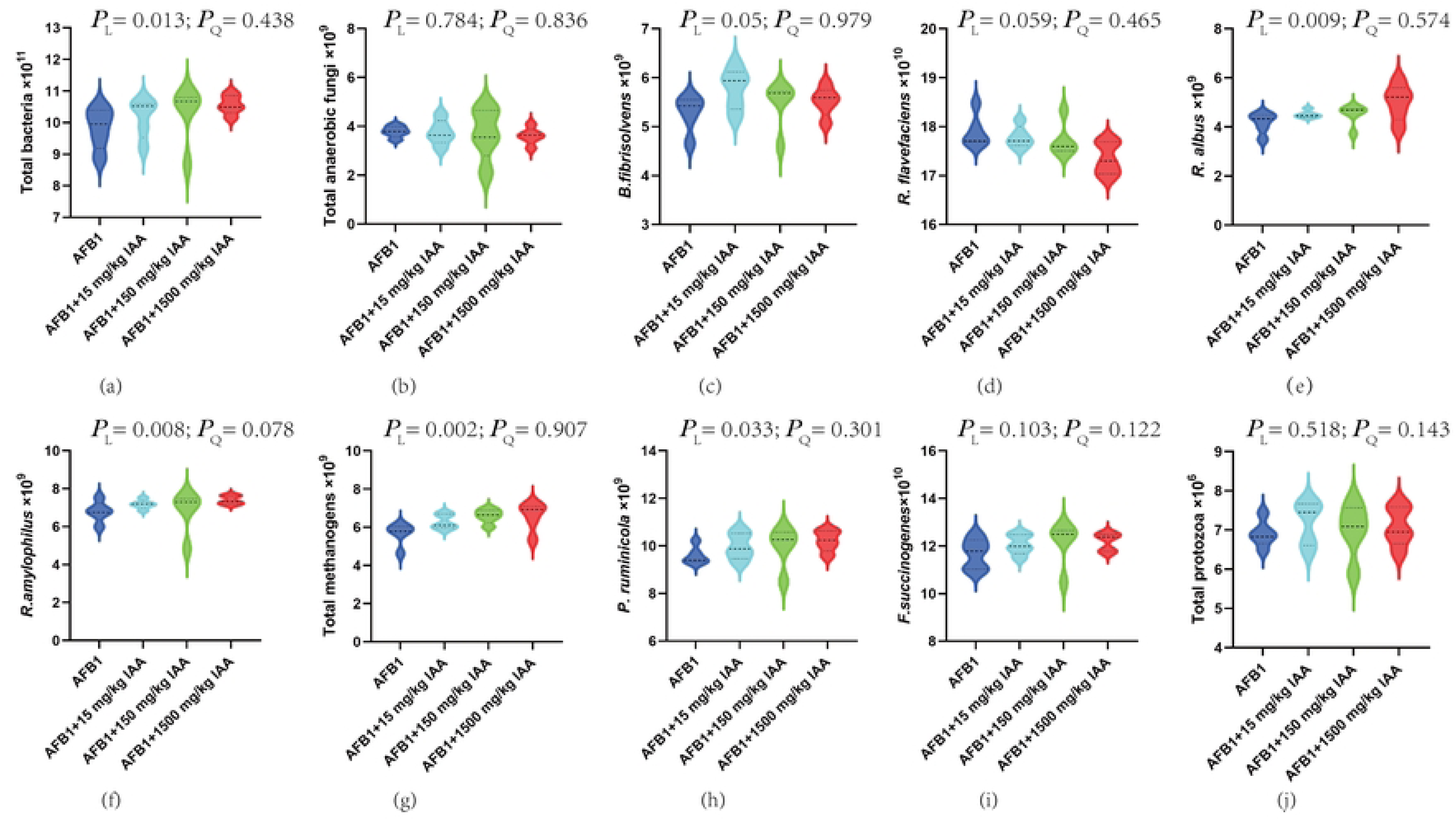
Effects of different concentrations of IAA on the activity of major fiber degrading enzymes in 24 h in vitro fermentation broth. PL represents a linear response, and PQ represents a quadratic linear response.

### 3.8 The impact of different concentrations of IAA on the microbial flora in 24-hour in vitro fermentation fluid

Figure 7 illustrates that when 15 and 150 mg/kg of IAA were added to the fermentation fluid, there was an increasing trend in the content of total bacteria and *Prevotella ruminicola*, showing significant linear effects with corresponding p-values of 0.013 and 0.033, respectively. Further research found that when the concentration of IAA was increased to 15, 150, and 1500 mg/kg, the content of *Butyrivibrio fibrisolvens, Megasphaera elsdenii, Ruminococcus amylophilus*, and total methanogenic archaea in the fermentation fluid also showed an upward trend. Similarly, this trend exhibited significant linear effects with corresponding p-values of 0.05, 0.009, 0.008, and 0.002, respectively. It is worth noting that *Ruminococcus amylophilus* not only showed a linear effect under the influence of changes in IAA concentration but also a quadratic effect, with the corresponding p-value being 0.078. This finding suggests that the variation in the content of *Ruminococcus amylophilus* may be influenced by a combination of factors, and its regulatory mechanism may be more complex.

**Fig.7.**
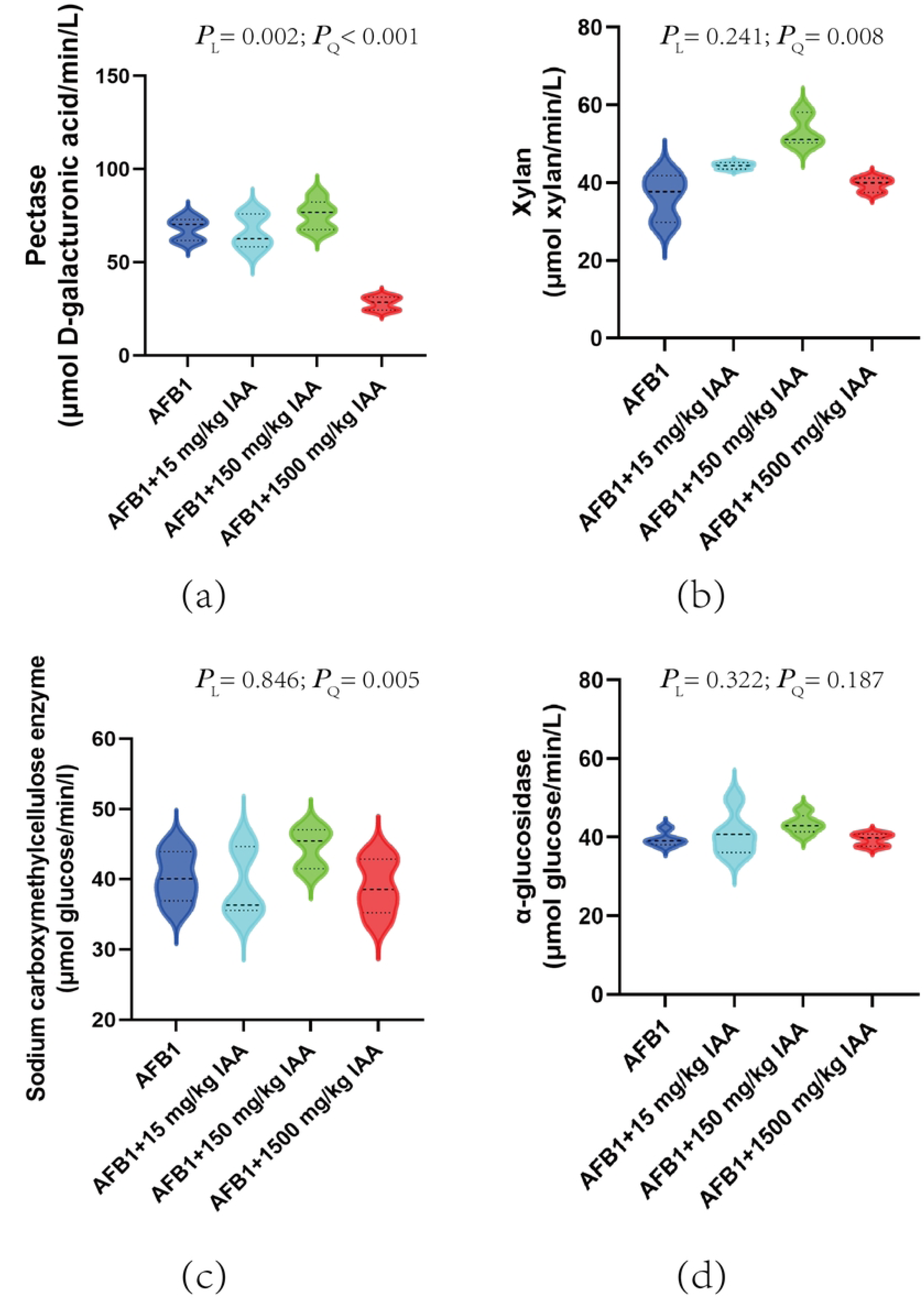
Effects of different concentrations of indole acetic acid on the microbial flora in the 24 h in vitro fermentation broth. P_L_ represents a linear response, and P_Q_ represents a quadratic linear response.

## 4. Discussion

Ruminant animals have a complex microbial community in their rumen that has been formed through long-term evolution. These microorganisms are interdependent and mutually restrictive. They are closely connected with the host’s immune system, nutritional metabolism, and interactions between organisms. Studies have found that certain microorganisms in the rumen have the ability to degrade AFB1. [26] The research by Upadhaya *et al.* [27] also confirmed that the microorganisms in the bovine rumen have the ability to degrade AFB1. This phenomenon can be attributed to several key factors: first, microorganisms reduce the toxicity of AFB1 through adsorption; second, enzymes secreted by rumen microorganisms can break down the structure of AFB1, converting it into other substances, further reducing its toxicity. During in vitro fermentation, 1 mg/kg of AFB1 was added to the fermentation fluid, and as the fermentation time increased, the concentration of AFB1 gradually decreased, reaching a stable state after a period of time. Especially when the fermentation time reached 48 h, the degradation efficiency of AFB1 can reach about 80%. In addition, research has shown that AFB1, as a harmful secondary metabolite widely present in feed and its raw materials, can disrupt the balance of microorganisms in the rumen when ingested by ruminants and enter the rumen, leading to changes in the protein content and pH value of the rumen microorganisms, thereby affecting the normal fermentation process of the rumen. [28, 29] Volatile fatty acids are the main source of energy for ruminants to obtain from feed [30], but the presence of AFB1 has a significant inhibitory effect on the rumen microbial community, leading to a decrease in the total production of volatile acids and a change in the fermentation type. In vitro fermentation experiments show that after exposure to AFB1, the abundance of core microorganisms in the rumen such as Prevotella and Fusobacterium decreased, affecting the balance and function of the microbial community. In addition, AFB1 also reduced the activity of xylanase, consistent with the changes of cellulolytic bacteria, while the activity of other enzymes such as pectinase did not change significantly, which may be related to the amount of AFB1 added. The rumen environment, especially the pH value, is crucial to the microbial balance and the health of the rumen [31]. The metabolic process in the rumen includes the degradation of cellulose and hemicellulose, the metabolism of soluble sugars, and the fermentation to produce organic acids and short-chain fatty acids. These findings indicate that AFB1 not only changes the fermentation pattern of the rumen but may also affect the nutritional absorption and health of ruminants.

IAA is an important biological regulatory substance, essential for the growth of forage, and is also considered to be an efficient and environmentally friendly additive with the potential to regulate the balance of animal intestinal microflora. Although research on IAA in rumen microorganisms is still in its infancy, this experiment has confirmed that adding different concentrations of IAA to rumen fluid containing AFB1 can effectively reduce the content of AFB1, especially when adding 7500 mg/kg IAA, the content of AFB1 significantly decreased. However, considering that high concentrations of IAA may affect the balance of the rumen, according to the “Safety Use Specifications for Feed Additives,” the recommended amount of tryptophan (the precursor of IAA) in the diet of ruminants is 0.1% (1 g/kg). In addition, some studies have shown that excessively high concentrations of IAA may change the animal intestinal microbial community [32]. Therefore, this experiment further explored the specific impact of IAA additions of 15 mg/kg, 150 mg/kg, and 1500 mg/kg on rumen fermentation parameters, aiming to find an appropriate level of IAA addition to alleviate the impact of AFB1 on the rumen fermentation process and improve animal health. This experiment studied the impact of different concentrations of IAA on VFAs, pH, and ammonia nitrogen production in in vitro fermentation fluid containing 1 mg/kg AFB1, and found that medium and low concentrations of IAA can significantly increase the content of total volatile fatty acids (TVFA), while high concentrations of IAA tend to decrease. The addition of IAA, especially medium and low concentrations, enhances the ability of the rumen microbial community to utilize carbon sources and stimulates the degradation of nutrients in the rumen. Acetic acid, propionic acid, and butyric acid are the main products of rumen fermentation and are crucial for energy metabolism and nutrient absorption [33-35]. In the experiment, after the addition of IAA, the content of acetic acid showed a linear upward trend, which is related to the activation of fiber-decomposing bacteria and enzyme activity by IAA, promoting the growth of fiber-decomposing bacteria and fungi. With the addition of IAA, the observed changes in the microbial community, such as Prevotella and Fusobacterium, are closely related to the adjustment of VFAs composition [36, 37]. These microorganisms decompose the fiber in the feed, promoting the production of acetic acid and propionic acid. Propionic acid is key in the gluconeogenesis process, affecting body fat and lactose synthesis, while butyric acid is easily absorbed by the rumen epithelial cells, providing energy [38-41]. The concentration of IAA has a significant regulatory effect on propionic acid production, with medium and low concentrations of IAA causing a linear increase in propionic acid content, while high concentrations of IAA reduce the level of propionic acid. The change in butyric acid content is also related to the amount of IAA added, with medium and low concentrations of IAA increasing butyric acid content, while high concentrations of IAA reduce butyric acid content. In addition, the regulation of the acetic acid/propionic acid ratio affects microbial protein synthesis and the structure of the rumen microbial community, which in turn relates to the digestion and nutritional metabolism of the whole body [42, 43]. In the rumen ecosystem, bacteria can degrade and utilize starch and plant cell wall polysaccharides, such as xylan and pectin, but they cannot degrade cellulose. These bacteria play an important role in the degradation of protein and the absorption and fermentation of peptides [44-50]. Finally, the addition of IAA has a significant impact on the changes in the rumen microbial community and the activity of the main fiber-degrading enzymes. Medium and low concentrations of IAA significantly increased the activity of pectinase, xylanase, and carboxymethyl cellulase, while high concentrations of IAA reduced the activity of these enzymes. These results show that IAA alleviates the imbalance of the rumen fermentation microecology caused by AFB1 by regulating the rumen microbial community and enzyme activity. In summary, the appropriate addition of IAA has a positive impact on the rumen microbial community and metabolic products, but high concentrations of IAA may inhibit the rumen fermentation process. These findings provide important information for optimizing rumen fermentation and improving the nutritional absorption of ruminants.

## 5. Conclusions

The addition of IAA has been proven to effectively enhance the ability of the native microbial community in the rumen to degrade AFB1, providing a new strategy to mitigate the potential threat of AFB1 to animal health. This study clarifies the key role of IAA in rumen fermentation, particularly showing significant effects in reducing the content of AFB1 within the rumen. After a comprehensive assessment of economic costs, the balance of animal gut microbiota, and health impacts, a recommended dosage of 15 mg/kg IAA is suggested to be added to feed. This dosage not only promotes the effective degradation of AFB1 in the rumen but also regulates the adverse effects of AFB1 on the rumen fermentation process, while ensuring the health and production performance of the animals.

## Supplementary Materials

The following supporting information can be downloaded.

## CRediT authorship contribution statement

LQ: Conceptualization.XY: Methodology. YZ: Software. LT: Validation. JS: Formal analysis, Writing - Original Draft. XM: Investigation. RZ: Resources. ZW: Data Curation. WH: Writing - Review & Editing. LC: Visualization. CW: Supervision. QL: Project administration.GG: Funding acquisition.

## Institutional Review Board Statement

The present study was approved by the Animal Health and Care Committee of the Shanxi Agricultural University (Shanxi, China) and conducted according to the Guidelines for the Experimental Animal Welfare of Ministry of Science Technology of China (approval code. SXAU-EAW-2022C.RD.010025174).

## Data Availability Statement

The data presented in this study are available on request from the corresponding author.

## Acknowledgments

The author thanks the National Natural Science Foundation of China for financially supporting our experiment.

